# The lack of peroxisomal Glycolate Oxidases 1 and 2 influences mitochondrial electron transport chain and its redox state under control and cadmium stress

**DOI:** 10.64898/2026.05.06.723131

**Authors:** Aurelio M. Collado-Arenal, María Rodríguez-Serrano, M. Ángeles Peláez-Vico, Laura C. Terrón-Camero, Felipe L. Pérez-Gordillo, Pablo Ranea-Robles, Luis Carlos López, Luisa M. Sandalio, María C. Romero-Puertas

**Author notes:** Education Faculty, University of Granada, Melilla, Spain (María Rodríguez-Serrano). Division of Plant Sciences and Technology, College of Agriculture Food and Natural Resources and Interdisciplinary Plant Group, University of Missouri, Columbia, MO 65211, USA. Bioinformatics Unit, Institute of Parasitology and Biomedicine “López-Neyra” (IPBLN-CSIC), 18016 Granada, Spain. Authors for correspondence: Dr. María C. Romero-Puertas, Dr. Luisa M. Sandalio.

## Abstract

The production of reactive oxygen species (ROS) in response to cadmium (Cd) has been extensively studied, demonstrating that they play a key role in the plant’s response to this heavy metal. While the role of enzymes like RBOHs has been thoroughly studied, the function of other ROS-producing enzymes, such as peroxisomal glycolate oxidase (GOX), remains largely overlooked. Peroxisomal GOX is a core metabolic enzyme of the photorespiratory pathway occurring in chloroplasts, mitochondria and peroxisomes. Using Arabidopsis (*Arabidopsis thaliana*) mutants lacking the main peroxisomal GOX genes, GOX1 (*gox1-1*) and GOX2 (*gox2-1*) we explored their function in plant response to Cd. Although photosynthetic capacity appears to be affected to the same extent in both mutants under control and Cd stress conditions, GOX2 seems to play a greater role in ROS production in response to the metal. Transcriptomic analyses on WT and *gox2-1* pointed to the mitochondrial electron transport chain (mETC) as a target of Cd stress. We further investigated the individual GOX1 and GOX2 functions in mETC regulation and redox state. Although oxidative ratio of mitochondria was higher in both mutants, it was more pronounced in the absence of GOX1. Furthermore, the mETC is affected in both mutants but the regulation of its components differs in each mutant. These results point out the different functions of the two photorespiratory GOX isoforms in Arabidopsis, leading to a better understanding of the photorespiratory pathway.

## INTRODUCTION

Although many natural chemical elements occur naturally within the Earth’s crust, including those that are essential to life, others such as cadmium (Cd) can be toxic if they accumulate. Cd is actually highly toxic even at low concentrations (Fu and Xi, 2020). While some geogenic processes are responsible for the accumulation of heavy metals in soil, human activities such as agriculture, industrial processes, electricity generation, mining, household waste and the most recent development of photovoltaic devices, electric vehicle batteries and wind turbines, contribute to environmental pollution (Food and Agriculture Organization of the United Nations, 2022; Okereafor et al., 2020; Yan et al., 2025). Therefore, heavy metals and in particular, Cd accumulation, are becoming a problem for agriculture, humans and the environment. Recently, it has been shown that around 15 % of cropland is affected by toxic metal pollution worldwide exceeding the agricultural threshold for at least one heavy metal. With a 9%, the global exceedance rate for Cd is the highest (Hou et al., 2025). In addition to policies to prevent an increase in soil contamination, research is needed to understand the effects of this heavy metal on plants and to prevent its entry into the food chain.

Although indirectly, one of the major effects of cadmium stress in plants is the production of reactive oxygen species (ROS), which occurs at low levels in the initial response, but at higher levels if the stress continues leading to oxidative stress damage (Cuypers et al., 2023; Sandalio et al., 2023). At low levels, ROS may act as signaling molecules regulating different processes in plant response to Cd such as photosynthesis efficiency, antioxidant activities, metal and ion homeostasis (Gupta et al., 2017; Hafsi et al., 2022). While different sources have been involved in plant ROS production after Cd treatment their contribution to plant responses to this heavy metal has not been deciphered so far (Sandalio et al., 2023). Although H_2_O_2_ may originate from plasma membrane NADPH oxidases (so-called RBOHs) in the early response to Cd (Garnier et al., 2006; Hafsi et al., 2022; Rodríguez-Serrano et al., 2016), different sources of ROS associated with other cell compartments may also be involved as different waves of ROS production have been described in different species (Garnier et al., 2006; Pérez-Chaca et al., 2014). Recently, a spatiotemporal distribution of redox changes by redox-sensitive green fluorescent proteins located in different compartments of the cell has been shown in plant response to Cd (Collado-Arenal et al., 2024). Peroxisomes are the organelles that undergo the most rapid changes in redox state in response to Cd (Collado-Arenal et al., 2024). In fact, peroxisomes are an important source of ROS with an essentially oxidative metabolism being key in maintaining the redox state of the cell (Molina-Moya et al., 2025). Different metabolic pathways such as β-oxidation, through Acyl-CoA oxidases (ACX) and photorespiration through glycolate oxidase (GOX), are involved in ROS production in these organelles. Polyamines catabolism, sarcosine oxidase (SOX), xanthine oxidoreductase (XOR) and urate oxidase (UO) are also involved in peroxisomal ROS production (Molina-Moya et al., 2025; Sandalio et al., 2023). In particular, photorespiration appears as a key pathway in the regulation of redox homeostasis in the cell (Foyer and Noctor, 2020). GOX, which is the primary peroxisomal enzyme in the photorespiration cycle, catalyzes oxidation of glycolate producing H_2_O_2_ during this reaction, being about the 70 % of the total H_2_O_2_ content in photosynthesizing C3 leaves (Noctor et al., 2002). GOX has been involved in plant responses to different stresses both biotic and abiotic (Kerchev et al., 2016; Rojas et al., 2012) and in particular, its activity has been shown to increase in plant response to Cd in different species such as pea, soybean, *Biscutella auriculata* and Arabidopsis (Gupta et al., 2017; Peco et al., 2020; Pérez-Chaca et al., 2014; Romero-Puertas et al., 1999). Five genes coding for GOX have been described in the Arabidopsis genome (Reumann et al., 2007) being *GOX1* and *GOX2* the major isoforms described in leaf peroxisomes (Rojas et al., 2012). Both genes are in the chromosome 3 of Arabidopsis located in tandem and it has been suggested that the duplication that gave rise to both genes is Brassicaceae specific (Kerchev et al., 2016). Individual *gox1* and *gox2* mutants showed similar GOX decreased activity and show no visible phenotype suggesting some redundancy in their function (Rojas et al., 2012). Recently, it has been described however that *gox1* mutant but not *gox2* decreased the photorespiratory phenotype of *cat2* mutants suggesting that both genes may also have different metabolic roles (Kerchev et al., 2016). We therefore investigated GOX-dependent plant responses to Cd by physiological, biochemical and transcriptomic approaches. Our results reveal similar and differential roles for *GOX1* and *GOX2* in plant responses to Cd suggesting new key function for these enzymes in maintaining efficient redox and energetic pathways and contributing to a better understanding of the photorespiratory pathway in plant responses to stress.

## MATERIALS AND METHODS

### Plant Material and cadmium treatment

All lines used in this study are in Columbia background (Col-0). Mutants *gox1-1* (SAIL_177_G11; *gox1*) and *gox2-1* (SALK_044052; *gox2*), were obtained from Nottingham Arabidopsis Stock Centre (NASC) and have been described before (Rojas et al., 2012). Arabidopsis lines expressing the genetically encoded biosensor GRX1-roGFP2 in mitochondria Mit-GRX1-roGFP2 was kindly provided by Dr A.J. Meyer (University of Bonn). The Mito-GRX1-roGFP2 transformants in the *gox1* and *gox2* backgrounds were selected, by using the GFP fluorescence as marker and genotype was confirmed by PCR (Kerchev et al., 2016). The seeds were disinfected on the surface and stratified for 24-48 hours at 4°C. They were then sown on Murashige and Skoog (MS) 0.5x solid medium with 3 % of sucrose (Murashige and Skoog, 1962). The seeds were grown at 22°C in 16 hours of light and 8 hours of darkness for 14 days (d). Seedlings were then transferred to plates with 0.5x liquid MS medium and grown for 24 h under these conditions. The medium was then supplemented or not with 100 µM CdCl_2_ for 30 min and 24 h. Samples for transcriptomic analysis and quantitative RT-PCR were then taken and frozen before use. The maximum quantum efficiency of photosystem II (Fv’/Fm’) was determined with a PAM-2000 chlorophyll fluorometer and the ImagingWinGigE software application (Walz; Effeltrich, Germany) on light-adapted plants. Arabidopsis plants for oxygen consumption rate (OCR) were grown hydroponically for 24 days in 0.5x Hoagland nutrient solution under long-day conditions at 20-22°C. Plants were either untreated or exposed to 50 µM Cd for 24 h prior to measurements.

### H_2_O_2_ quantification and enzymatic activities

Hydrogen peroxide (H_2_O_2_) was quantified by spectrofluorimetry as described before (Romero-Puertas et al., 2004), using horseradish peroxidase and homovanillic acid (excitation: 315 nm; emission: 425 nm) in 50 mM HEPES pH 7.5. Commercial H_2_O_2_ was used for the standard curve with known concentrations to quantify samples. For enzymatic activities whole seedlings were homogenized as described by Rodríguez-Serrano et al. (2016) and catalase (CAT; EC 1.11.1.6) and GOX activities were measured according to Aebi (1984) and Kerr and Groves (1975), respectively. Proteins were quantified using Bradford Protein Assay (Bio-Rad) and BSA (bovine serum albumin (BSA) was used for the standard curve.

### RNA isolation and RT-PCR analysis

Total RNA was isolated from seedlings using the TRIzol reagent ® (MRC) according to the manufacturer’s instructions. RNA was reverse transcribed using the PrimeScript RT Master Mix (Takara) following the instructions of the commercial company. In brief, each 20 µl reaction contained either 1 µl cDNA or a dilution, 200 nM of primers each, and 1x TB Green Premix Ex Taq (Takara). Quantitative real-time PCR was performed on an iCycler iQ5 (Bio-Rad) as described before (Terrón-Camero et al., 2020). Relative expression of genes was normalized using *TUB4* that was selected for normalization by the GrayNorm algorithm (Remans et al., 2014) from five candidate reference genes as we described previously in (Terrón-Camero et al., 2020). Results were calculated with the ratio according to the Pfaffl method (Pfaffl, 2001). Primers used are described in Suppl. Table S1.

### Bioinformatics

WT and *gox2* transcriptomic data under control and Cd treatment are deposited in the Gene Expression Omnibus repository (GSE199325; Terrón-Camero et al., 2022). Functional enrichment analyses were performed on differentially expressed gene sets from *Arabidopsis thaliana* WT and *gox2* under control and Cd stress (30 min. and 24h). Gene identifiers were based on TAIR annotations, and all genes detected in the experiment were used as the reference universe. The non-overlapping genes between the data sets of interest were computed using the Venny algorithm (http://bioinfogp.cnb.csic.es/tools/venny/). Gene Ontology (GO) enrichment analyses for the categories of biological process, molecular function, and cellular component were performed using the R clusterProfiler package, together with annotation resources from org.At.tair.db and AnnotationDbi. Multiple testing correction was applied using the Benjamini-Hochberg method. Enrichment results were visualised using ggplot2, enrichplot, and corrplot, including bar charts, dot plots, and GO plots. Expression patterns of genes associated with significantly enriched functional categories were summarised using heatmaps, and the overall distribution of differential expression was represented using volcano plots, both generated following enrichment analysis. All analyses and visualisations were performed in R, using additional packages such as edgeR, dplyr, tidyr, ggrepel, and RColorBrewer.

We used the ROSMETER platform to identify transcriptomic tracks in plant responses to Cd on the basis of ROS type and origin (Rosenwasser et al., 2013). Significantly enriched (*P* < 0.05) GO terms from data sets of interest were analysed using Mapman software (https://mapman.gabipd.org/), which displays large datasets in diagrams of metabolic pathways.

### Mitochondrial respiration rate

Mitochondrial respiration was assessed by measuring the Oxygen Consumption Rate (OCR) using an Agilent Seahorse XFe24 Extracellular Flux Analyzer. Leaf discs (5 mm diameter) were excised and immobilized in Seahorse XFe24 cell culture microplates using 5% Leukosan® adhesive in 0.25% Plant-Agar (Sew et al., 2013). Discs were maintained in assay medium containing 30 mM MES and 0.2 mM CaCl₂, adjusted to pH 5.6. Prior to measurements, plates were incubated for 1 h at 22°C in the dark to allow equilibration. OCR was then monitored with measurement cycles performed every 10 min. After an initial stabilization period (45 min), sequential injections were performed as follows: 2 mM salicylhydroxamic acid (SHAM), 0.08 mM rotenone, and 0.1 mM KCN at 45, 105, and 165 min, respectively (Hu et al., 2016; Maliandi et al., 2015). These inhibitors were used to dissect mitochondrial respiratory pathways, including alternative oxidase (AOX) and cytochrome pathway contributions. Each condition included six biological replicates.

### Mitochondrial redox state analysis

GRX1-roGFP2 oxidation rate located in seedlings from WT, *gox1* and *gox2* mitochondria was monitored by sequential excitation at 405 ± 8nm (Ex405) and 480 ± 8nm (Ex488) and the emission was recorded at 530 ± 10nm (Em530) in a BLACK SENSOPLATE 24 well (Greiner Bio-One 662892) using a CLARIOstar plate reader (BMGLabtech, Ortenberg, Germany) as described elsewhere (Collado-Arenal et al., 2024).

### Statistical analysis

Mean values for the quantitative experiments described above were obtained from at least three independent experiments, with no less than three independent samples per experiment. Statistical analyses were performed using a one- or two-way ANOVA test, when necessary, followed by a student’s t-test (p-value < 0.05) or Tukey multiple comparison test (p-value < 0.05), respectively. The analyses were carried out with the aid of IBM SPPS Statistics 24 and GraphPad Prism 6. Error bars representing standard error (SEM) are shown in the figures.

## RESULTS

### GOX is involved in ROS production following Cd treatment in Arabidopsis

H_2_O_2_ production has previously been found to increase after Cd treatment in Arabidopsis seedlings and leaves, with some evidence suggesting that GOX, the enzyme catalysing the first step of photorespiration, may be an important source of Cd-induced H_2_O_2_ (Gupta et al., 2017; Rodríguez-Serrano et al., 2016). We firstly determined, in control and Cd-treated WT plants, H_2_O_2_ production in Arabidopsis seedlings treated with Cd at different time-points (30 min and 24 h). Similar to previously reported, following 30 minutes of Cd treatment, a minor rise in H_2_O_2_ was detected, however this increase was not found to be significant (Fig. 1A). A 24-hour exposure to Cd resulted however in a significant increase in H_2_O_2_ production as illustrated in Fig. 1A. We then analysed GOX activity in WT seedlings treated with Cd and observed a significant increase after 24 h (Fig. 1B). To determine whether the increased GOX activity was related to either of the two main GOX-coding genes, we used the *gox1-1* and *gox2-1* mutants (*gox1* and *gox2* from now on), which are affected in *GOX1* and *GOX2* genes, respectively (Rojas et al., 2012). We observed a significant increase of GOX activity in *gox1* but not in *gox2* seedlings (Fig. 1B), suggesting that *GOX2* may be the main gene involved in the response to Cd treatment. Both mutants showed a residual GOX activity of about 53 % and 65% of the WT for *gox1* and *gox2* respectively (Fig. 1B). We did not observe an increase in H_2_O_2_ content after 24 h of Cd treatment in any of the *gox* mutant lines however (Fig. 1 A). Catalase is the main antioxidant in peroxisomes involved in H_2_O_2_ removal associated to photorespiration (Mhamdi et al., 2012). Therefore, we also analysed the catalase activity of Cd-treated seedlings and found that it is increased significantly only in the *gox1* mutants (Fig. 1C). This result suggests that CAT may be involved in the removal of H_2_O_2_ in the *gox1* response to 24 h of Cd treatment. We also measured as an indicator of stress, the maximum quantum efficiency of Photosystem II under light-adapted conditions in WT, *gox1* and *gox2* under control and Cd stress for 24 and 72 h (Fig. 1D). Under Cd stress, all lines display a decline of PSII maximum efficiency (Fv’/Fm’), which is time-dependent. Both mutants, *gox1* and *gox2* showed a significant lower PSII maximum efficiency than WT under control and Cd stress (Fig. 1D).

**Figure 1:**
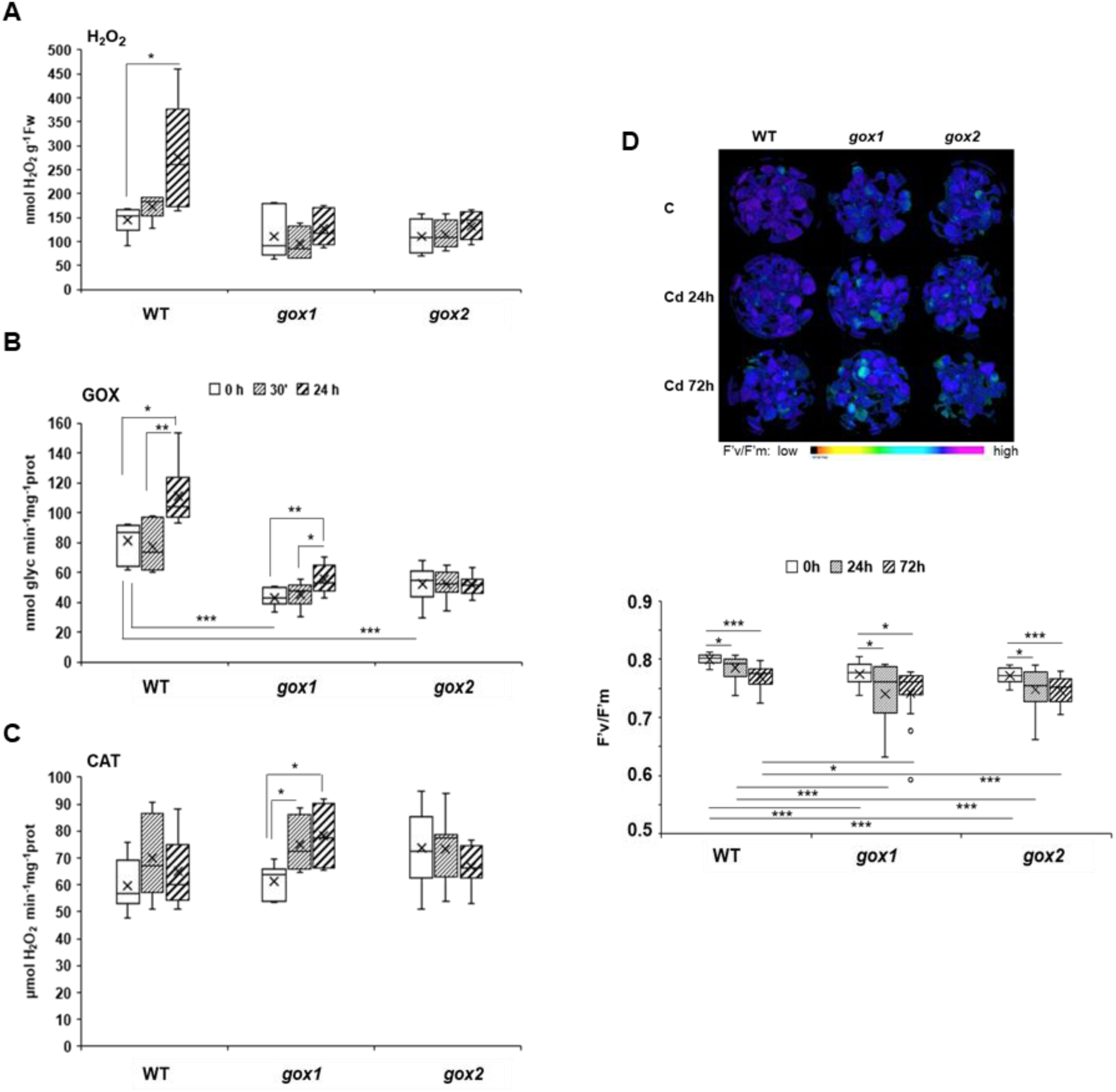
ROS production and peroxisomal antioxidant activity after Cd treatment. A) H_2_O_2_ content assayed by fluorimetry in acid extracts from WT, *gox1 and gox2* seedlings after Cd treatment (30 min-24 h). B) GOX and C) CAT activities in WT, *gox1 and gox2* seedlings extracts after Cd treatment (30 min-24 h). D) Color-coded image of PSII maximum efficiency (Fv’/Fm’) of WT, *gox1 and gox2* seedlings after Cd treatment (24 h-72 h) and boxplot showing Fv’/Fm’ values under these conditions. Values are means ± SE of at least three experiments with three independent biological replicates each. Asterisks denote significant differences between data according to the Student’s t-test (p-value < 0.05). The absence of an asterisk denotes no significant differences.

### Cd-dependent transcriptome footprint is related to different ROS signatures

Although the Cd effect in plants has been partly attributed to ROS (Cuypers et al., 2023; Molina-Moya et al., 2025), little is known about the underlying molecular mechanisms of ROS-dependent regulation. Furthermore, the increase of GOX activity in WT seedlings indicates that this enzyme may play a role in plant responses to Cd treatment. Considering the GOX activity in *gox1* and *gox2* mutants, we conducted a time-course transcriptomic analysis of wild-type (WT) and *gox2* Arabidopsis seedlings treated with cadmium (Cd) to identify genes that were differentially regulated in both lines in response to the heavy metal.

We analysed the transcriptomes at 30 min after Cd treatment, in parallel to the early slight increase of H_2_O_2_ in WT plants, and at 24 h post-treatment, when GOX activity is induced and the increase of H_2_O_2_ is significant. We then used the ROSMETER bioinformatics platform to investigate the occurrence of ROS-related transcriptomic signatures in our transcriptomic changes, relating them to ROS type and origin (Rosenwasser et al., 2013). Significantly, the highest transcriptome correlation values after 30 min of Cd treatment (Fig. 2A) were found in relation to the changes observed in the transcriptome after 12 h rotenone treatment, which affects the mitochondrial Complex I (Garmier et al., 2008); the transcriptome changes of *flu* mutants due to early singlet oxygen production because of the accumulation of chlorophyll precursors (op den Camp et al., 2003); and the transcriptome changes after ozone treatment, related with the initial apoplastic ROS production and amplification by RBOHs (Wrzaczek et al., 2010). Interestingly, a high negative correlation was found between early Cd-dependent transcriptomic changes and the changes observed in antisense AOX1 plants and CAT-deficient mutants related with mitochondria and peroxisomes oxidative metabolism, respectively (Fig. 2A).

**Figure 2:**
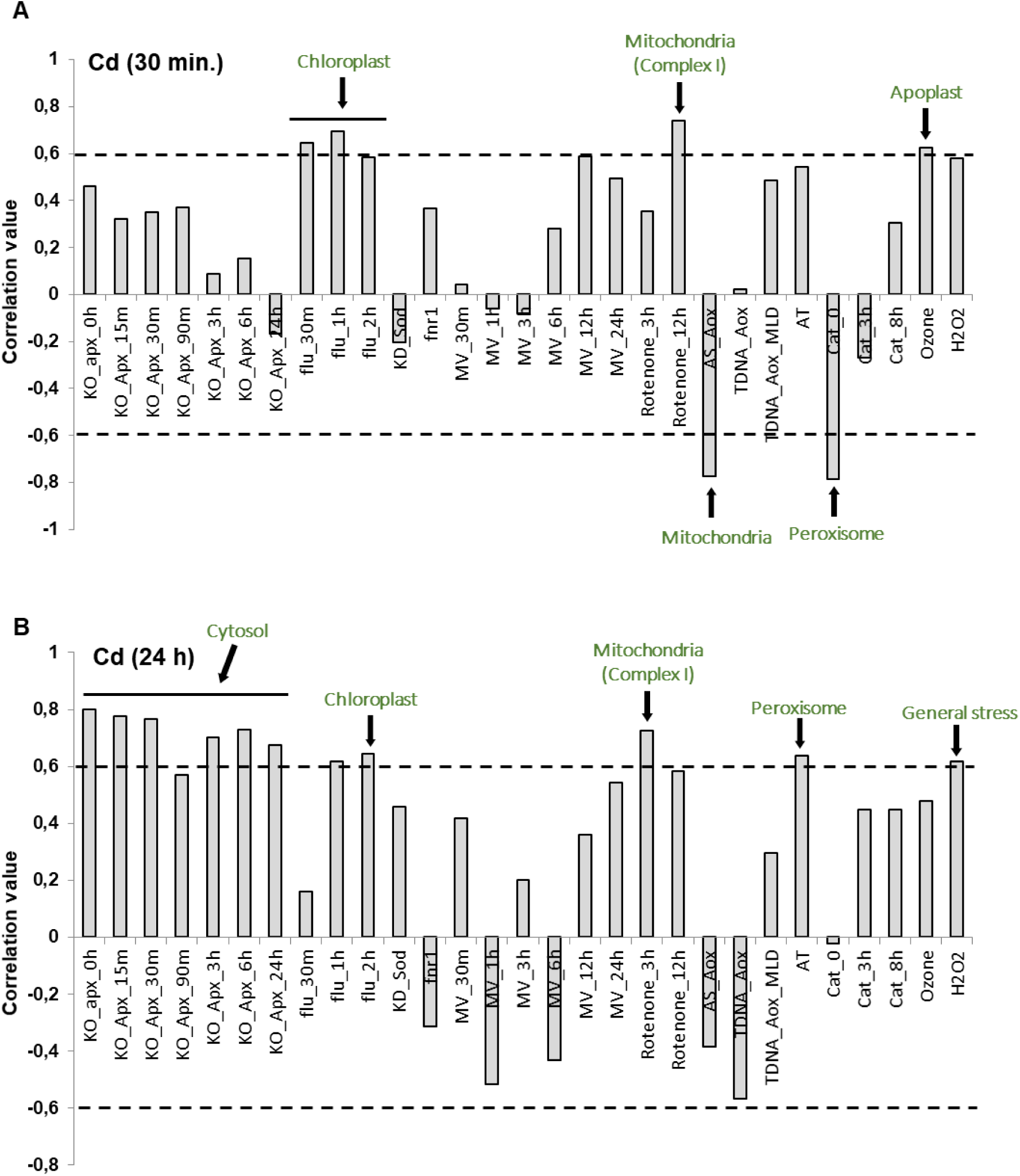
Analysis of Cd transcriptome using the ROSMETER platform. Correlation values of changes in the transcriptome in WT Arabidopsis plants treated with Cd for 30 min (A) and 24 h (B) generated by the ROSMETER platform. Correlation values (*y* axis ordinate) were obtained as described by Rosenwasser et al. (2011, 2013) from the Cd transcriptome and transcriptomes of the individual ROS-producing treatments compiled in the ROSMETER platform (*x* axis abscissa) detailed in Rosenwasser et al. (2013). “1” indicates complete correlation and “0” indicates no correlation. Positive and negative data correspond to positive and negative correlation, respectively, between the transcriptomes. Correlation values above 0.4 can be considered significant correlation values that provide biological insights (Rosenwasser *et al.,* 2013). Higher correlation values (arrows) at 30 min. relate to mitochondrial stress (Rotenone treatment), apoplast stress (ozone treatment) and chloroplast stress (flu mutants). Higher correlation values (arrows) at 24 h include also peroxisomal stress (AT, aminotriazol treatment), general stress (H_2_O_2_ treatment), and cytosol stress (*apx1* mutants exposed to high light).

Changes in the transcriptome after 24 h of Cd treatment and changes due to rotenone treatment and singlet oxygen production in the *flu* mutant maintained a significant correlation (Fig. 2B). Additionally, correlations with changes in ROS associated with the cytosol and peroxisomes have been added at this time-point (Fig. 2B).

### GOX2-dependent genes in early plant responses to Cd

Transcriptome analysis provides a deeper insight into both WT and *gox2* mutant by identifying peroxisomal-dependent genes that regulate plant responses to the heavy metal. Surprisingly, most of the genes regulated by Cd in WT are not regulated in *gox2* mutants leading to a high percentage of GOX2-dependent genes in plant responses to Cd (Fig. 3). About 500 transcripts were significantly regulated in early WT response to Cd, being about 75 % of them up-regulated (Fig. 3A). However, about 800 transcripts were regulated in *gox2* mutants, being in a similar number the up- and down-regulated (Fig. 3A). In total, we found 463 differential expressed genes (DEGs) regulated in WT but not in *gox2* mutants after 30 min Cd treatment, which we call GOX2-dependent genes (Suppl. Table S2). A significant enrichment in GO functional categories from Biological Processes related to hypoxia, fatty acid and plant type cell wall modification were found among early GOX2-dependent regulated transcripts (Fig. 3 B; Suppl. Table S3). Different activities are enriched also within the Molecular Function categories, such as dioxygenase, transaminase and transferase activity (Suppl. Table S3; Suppl. Fig. S1). Interestingly, the CCR4-NOT core complex, involved in regulating gene expression at different level, including transcription, translation, and mRNA degradation; and the plasma membrane are the main GOs related with Cellular Components at this early stage of plant response to Cd (Suppl. Table S3; Suppl. Fig. S1A).

**Fig. 3:**
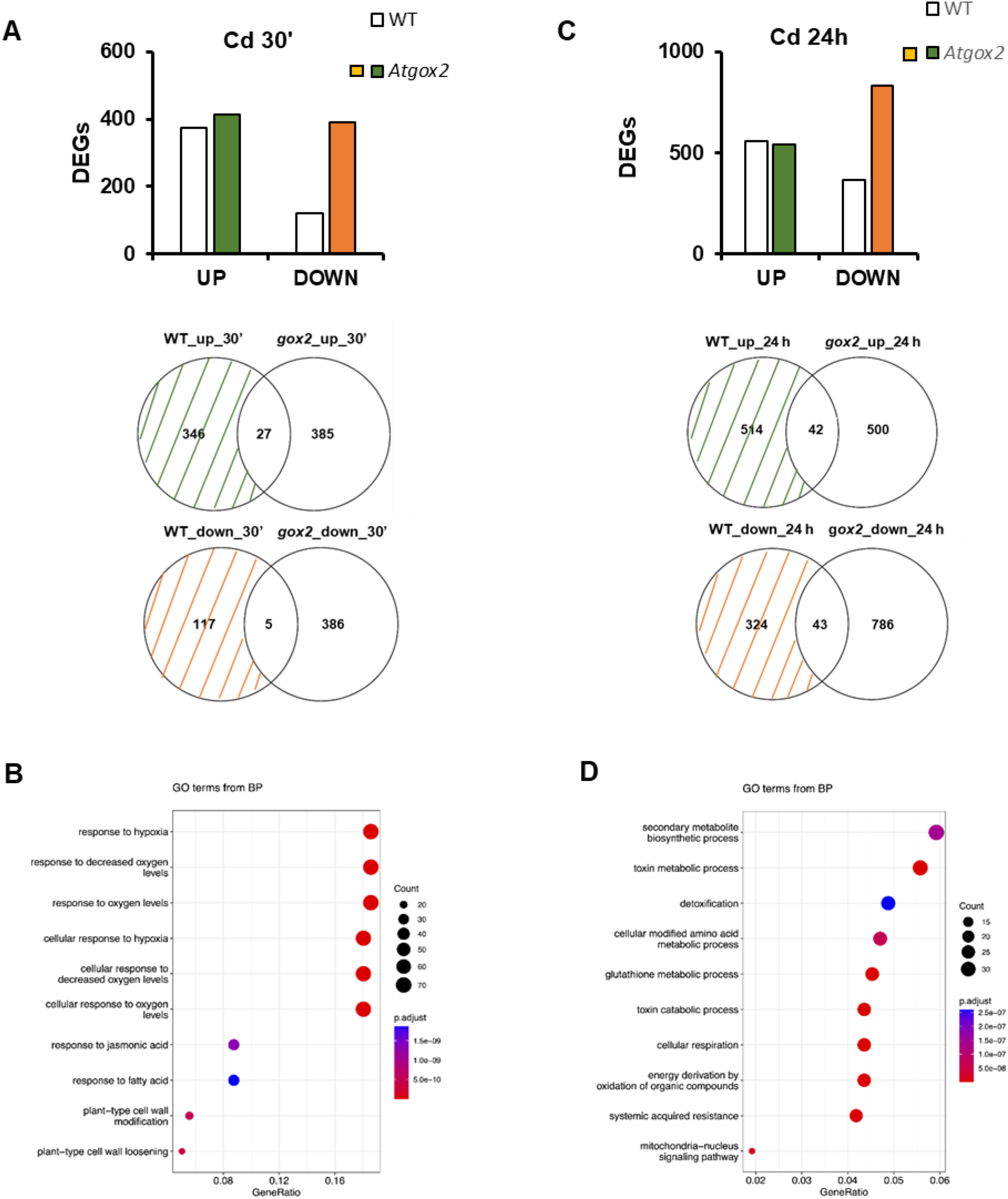
Changes in global transcript expression in the *gox2* mutant compared to wild type (WT) in response to Cd treatment. Number of up- and down-regulated genes in WT and *gox2* mutants after 30 min (A) and 24 h (C) of Cd treatment, and Venn diagrams showing the overlap between gene expression changes in WT and *gox2* mutants. Transcript expression altered in WT seedlings, but not *gox2* (GOX2-dependent) is marked by green (up-regulated) and orange (down-regulated) coloured stripes. Main Biological Process categories after gene ontology (GO) enrichment of GOX2-dependent genes at 30 min (B) and 24 h (D) after Cd treatment (Suppl. Tables S3 and S4). Normed to frequency of class over all ID numbers on *x* axes.

### GOX2-dependent genes in late plant responses to Cd

After 24 h Cd treatment, about 950 transcripts were significantly regulated in later WT response to Cd, being about 60 % of them up-regulated (Fig. 3C). However, about 1500 transcripts were regulated in *gox2* mutants, being the percentage of up-regulated transcripts similar to the WT but almost double of the down-regulated genes (Fig. 3C). In total, we found around 840 DEGs regulated in WT but not in *gox2* mutants after 24 h Cd treatment, which we call late GOX2-dependent genes (Suppl. Table S2). A significant enrichment in Biological Processes-GO functional categories related to toxin metabolic processes, secondary metabolites synthesis, glutathione metabolic processes and mitochondrial-nucleus signaling pathway among others are significantly overrepresented (Fig. 3D; Suppl. Table S4). Different activities are enriched within the Molecular Function categories, such as NADH dehydrogenase, oxidoreductase and glutathione-S-transferase (GST) among others (Suppl. Table S4; Suppl. Fig. S1B).

### GOX2 and Cd-dependent genes involved in ROS and redox metabolism

To check whether GOX2-dependent genes in plant response to Cd were related with ROS and/or redox metabolism we examine those genes in depth and found significantly represented the categories “removal of superoxide radicals and its regulation” and “GST activity” and “metabolic process” after 30 min. treatment (Fig. 4; Suppl. Table S5); and “GST activity” and “metabolic process” after 24 h treatment (Fig. 4; Suppl. Table S5). All genes in these categories are induced, with the exception of the gene involved in the regulation of the removal of superoxide radicals, which is repressed (Fig. 4).

**Figure 4:**
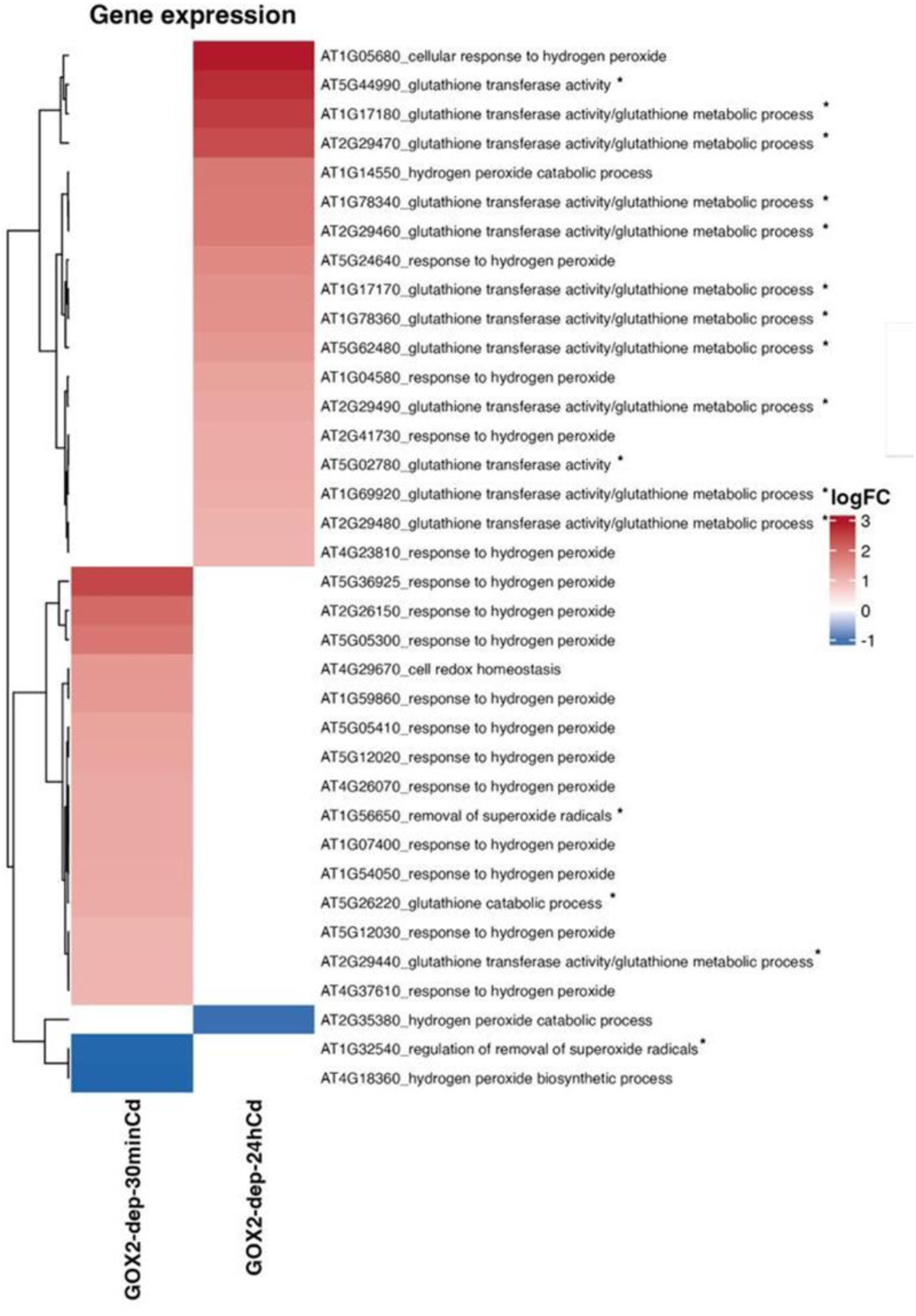
Analysis of GOX2-dependent genes in plant response to Cd related with ROS/GSH/ASC and redox metabolism. Heatmap of the GOX2-dependent differentially regulated genes (p-value < 0.05) in plant response to Cd, within the ROS/GSH/ASC and redox-related GO categories. Significantly over-represented functional categories are highlighted by asterisks (Suppl. Table S5). Locus identifiers (Suppl. Table S5) have been accompanied by Gene Ontology category related to oxidative metabolism. Glutathione transferase activity/glutathione metabolic process:GO:0004364/GO:0006749, response to hydrogen peroxide:GO:0042542, regulation of removal of superoxide radicals:GO:2000121, glutathione transferase activity/glutathione peroxidase activity:GO:0004364/GO:0004602, regulation of hydrogen peroxide metabolic process:GO:0010310, response to hydrogen peroxide/response to redox state:GO:0042542/GO:0051775, cell redox homeostasis:GO:0045454, positive regulation of hydrogen peroxide biosynthetic process:GO:0010729, hydrogen peroxide catabolic process:GO:0042744, hydrogen peroxide biosynthetic process:GO:0050665, cellular response to hydrogen peroxide:GO:0070301, glutathione peroxidase activity:GO:0004602, monodehydroascorbate reductase (NADH) activity:GO:0016656.

### Cd treatment triggered a GOX-dependent regulation of mitochondrial ETC

Interestingly, the main organelle within the Cell Components in the GOX2-dependent DEGs in plant response to Cd 24 h is the mitochondria (Fig. 5 A; Suppl. Table S4; Suppl. Fig. S1B). In fact, most of the mitochondrial DEGs are induced after 24 h Cd treatment being Complex I the most affected and in a lesser extent Complex IV as it can be observed in the image adapted from Map-man software (Fig. 5B; Table 1). Transcriptomic data shows that *AOX1* gene is also up-regulated after 24 h Cd treatment (Table 1). We then analysed by qRT-PCR the expression of some of the genes related with the mitochondrial ETC (Table 1; Fig. 5C). Most of the genes analysed were up-regulated after 24 h Cd treatment in WT seedlings but not in *gox2* mutants verifying transcriptomic analysis. *AOX1* however was also up-regulated by the treatment in *gox2* background although in a lesser extent than in WT (Fig. 5C). Furthermore, *AOX1* was already slight but significantly induced under control conditions in *gox2* compared with WT (Fig. 5C). *CyB* gene was also up-regulated in *gox2* compared with WT under control conditions (Fig. 5C). At early timepoint, only *CyB* was regulated in WT response to Cd being down-regulated in this case (Suppl. Fig. S2).

**Figure 5:**
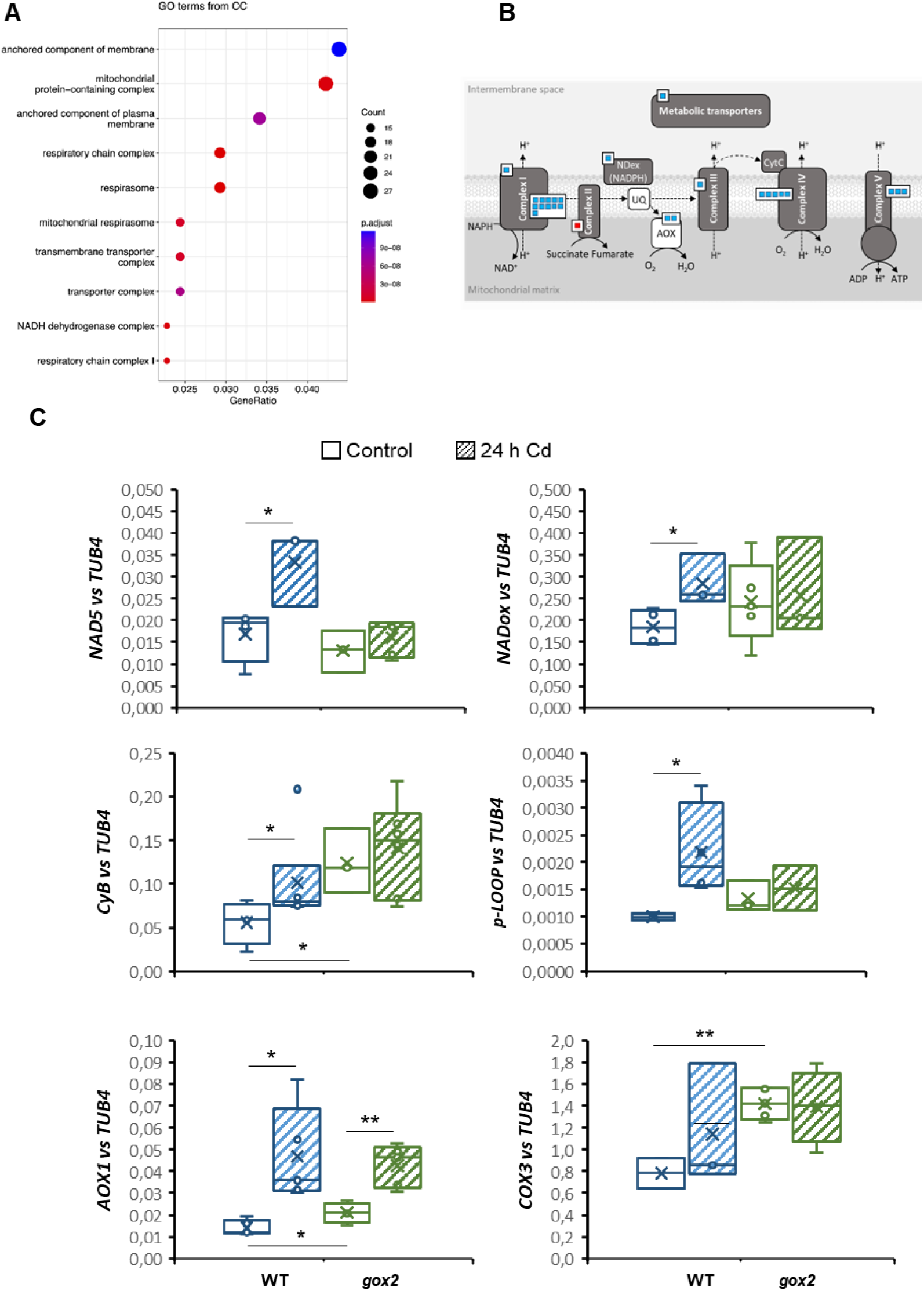
GOX2-dependent genes and transcriptomic changes by Cd in mitochondrial ETC. A) Main Cellular Component categories after gene ontology (GO) enrichment of GOX2-dependent genes at 24 h after Cd treatment (Suppl. Table S2). Normed to frequency of class over all ID numbers on *x* axes. B) Modified Mapman graph localizing GOX2-dependent DEGs related with the mitochondrial ETC. C) Quantitative real-time PCR analyses of genes related to ETC (Table 1). Each gene was normalized against *TUB4* expression. Primers used are described in Suppl. Table S1. Asterisks denote significant differences between data according to the Student’s t-test (p-value < 0.05). The absence of an asterisk denotes no significant differences.

**Table 1:** GOX2-dependent genes regulated by Cd after 24 h treatment related with mitochondrial electron transport / ATP synthesis following Map-man analyses.

| ID | Gene | Function | Length<br>(aa) | Molecular<br>weight | Isoelectric<br>point |
| --- | --- | --- | --- | --- | --- |
| AT2G07689 | NAD-OX | NADH-ubiquinone/plastoquinone Complex I | 214 | 22738,6 | 10,13 |
| AT2G07711 | NAD5 | NADH dehydrogenase 5B |  |  |  |
| AT3G22370 | AOX1 | Alternative oxidase 1A | 354 | 39979,6 | 8,71 |
| AT2G07727 | CYB | cytochrome b | 393 | 44146,4 | 7,29 |
| AT2G07687 | COX3 | cytochrome c oxidase subunit 3 | 265 | 29742,4 | 6,89 |
| AT4G05340 | P-LOOP | P-loop containing nucleoside triphosphate hydrolases superfamily protein | 96 | 11188,8 | 9,07 |
| ATMG00650 | NAD4L | NADH dehydrogenase subunit 4L | 100 | 11135 | 6,67 |
| ATMG01320 | NAD2B | mitochondrial NAD(P)H dehydrogenase | 306 | 35283,6 | 8,55 |
| ATMG01120 | NAD1B | mitochondrial NAD(P)H dehydrogenase | 91 | 9965,5 | 4,58 |
| ATMG00070 | NAD9 | mitochondrial NAD(P)H dehydrogenase | 190 | 22687,4 | 8,2 |
| ATMG00510 | NAD7 | mitochondrial NAD(P)H dehydrogenase | 394 | 44577,4 | 7,88 |
| AT2G07785 |  | NADH-ubiquinone oxidoreductase putative | 99 | 10610,4 | 10,1 |
| ATMG00513 | NAD5A | mitochondrial NAD(P)H dehydrogenase | 482 | 52456,9 | 7,73 |
| ATMG00270 | NAD6 | mitochondrial NAD(P)H dehydrogenase | 205 | 23493,8 | 10,24 |
| ATMG00060 | NAD5C | mitochondrial NAD(P)H dehydrogenase | 180 | 19353,9 | 9,43 |
| ATMG00285 | NAD2A | mitochondrial NAD(P)H dehydrogenase | 192 | 21494,1 | 6,93 |
| AT2G20800 | NDB4 | mitochondrial NAD(P)H dehydrogenase | 582 | 65371,2 | 9,32 |
| ATMG00160 | COX2 | cytochrome c oxidase subunit 2 | 260 | 29375,6 | 5,51 |
| ATMG01360 | COX1 | cytochrome c oxidase subunit 1 | 527 | 57995,9 | 8,05 |
| ATMG00730 | COX3 | cytochrome c oxidase subunit 3 | 265 | 29742,4 | 6,89 |
| ATMG00410 | ATP6-1 | ATPase subunit 6 | 385 | 42351,9 | 4,41 |
| ATMG00640 | ORF-25 | b subunit of mitochondrial ATP synthase | 192 | 21570,1 | 9,88 |
| ATMG01170 | ATP6-2 | ATPase subunit 6 | 349 | 39738,9 | 6,84 |

### Mitochondrial respiration is affected by Cd treatment and regulated by GOX

In the light of the transcriptomic results obtained on the regulation of the mitochondrial electron transport chain genes by Cd and because they are differentially regulated in *gox2* mutants, we decided to analyse respiration in WT, *gox2* and *gox1* seedlings to ascertain the possible regulation of respiration by the two main isoforms of GOX in photosynthetic tissues under control and Cd stress. We observed a significant induction of more than double of the mitochondrial respiration under Cd stress compared with control conditions in WT. Treatments with different inhibitors affecting AOX (SHAM), Complex I (Rotenone) and Complex IV (KCN) suggest that after Cd treatment an inhibition of the Complex I is apparently occurring and the Cd-dependent increase may be due to an induction of the Complex IV and AOX1 (Fig. 6A; Suppl. Table S6; Suppl. Fig. S3). Interestingly, both mutants *gox1* and *gox2* showed a higher respiration rate than WT seedlings under control conditions being 2.3 and 2.7 times greater in *gox1* and *gox2* respectively (Fig. 6A). However, no significant induction was observed in the mutants lacking *GOX1* or *GOX2* due to Cd treatment (Fig. 6A). Nevertheless, the contribution of AOX, Complex I, and complex IV to respiration differed in control and in response to Cd treatment in the mutant lines compared with WT. Thus, the increase of respiration observed in *gox1* lines under control conditions was mainly due to a higher AOX activity and in less extent to a slight increase of Complex I and Complex IV function. Under Cd treatment however, the increase observed in *gox1* lines is mainly due to an induction of the Complex I and AOX activity (Fig. 6A). The increase observed in *gox2* lines under control conditions compared to the WT was mainly due to the induction of Complex I and in less extent to an increase in AOX activity (Fig. 6). Although Cd treatment does not increase mitochondrial ETC in *gox2* lines, differences in the contribution of the different components were observed, with AOX having the main contribution, while an inhibition of the Complex I is taking place (Fig. 6A). All these results together suggest a deregulation of mitochondrial respiration in *gox*’s mutants under control and Cd treated seedlings, affecting differentially to the components of the ETC depending on the lack of *GOX1* or *GOX2*.

**Figure 6:**
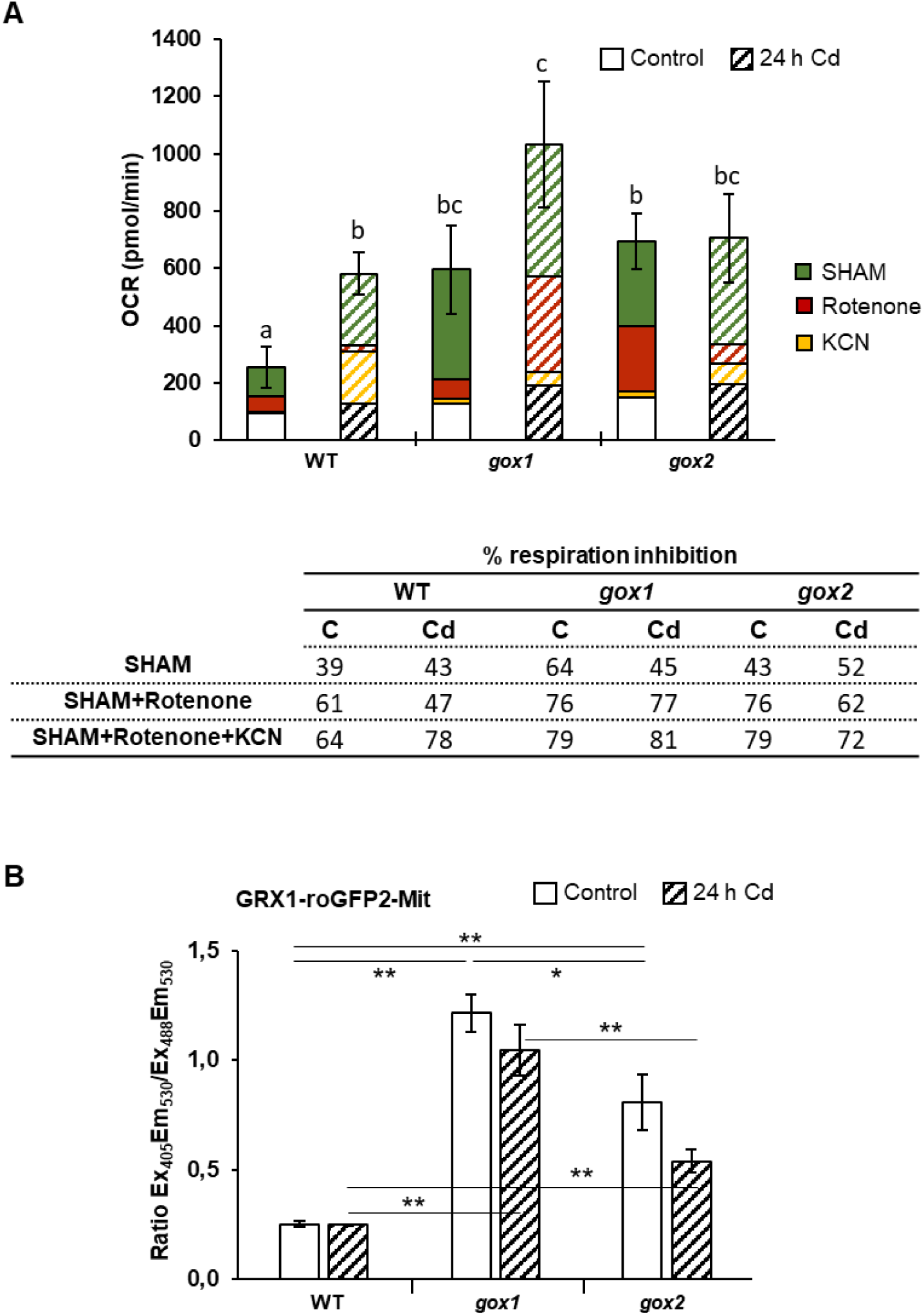
Cadmium effect on mitochondrial respiration and redox state in WT and *gox* mutants. A) Mitochondrial respiration in WT and *gox1* and *gox2* mutants treated or not with Cd for 24 h. The effect of complex I inhibitor, rotenone; the alternative oxidase inhibitor, SHAM and the complex IV inhibitor, KCN were used also and the percentage of respiration due to the different components was calculated. B) Redox state of WT and *gox1* and *gox2* mitochondria under control and Cd treatment (24 h), measured by ro-GRX1-GFP2 targeted to the organelle. Asterisks denote significant differences between data according to the Student’s t-test (p-value < 0.05). The absence of an asterisk denotes no significant differences.

### Redox state of mitochondria is affected by photorespiratory GOX

As the absence of the main GOX isoforms affects mitochondrial ETC, which is an important source of reactive oxygen species in the cell, we studied the redox state of mitochondria under control conditions and in response to Cd in the WT and *gox1* and *gox2* lines. To this end, we obtained double mutants containing the GRX1-roGFP2 biosensor localized in the mitochondria in *gox*’s lines. We do not observed changes in the oxidation state of mitochondria in WT Arabidopsis seedlings treated with Cd for 24 h suggesting that the redox state of this organelles is not affected at this time-point (Fig. 6B). Interestingly, we found that mitochondria from both, *gox1* and *gox2* lines, showed a higher 405/488 nm fluorescence ratio than WT suggesting that under control conditions these organelles are more oxidized due to the absence of GOX’s being mitochondria from *gox1* lines the most affected (Fig. 6B). No significant changes were observed either in the mutants treated with Cd compared with their control conditions although their 405/488 nm fluorescence ratio was significantly higher than in the WT also under this condition (Fig. 6B).

## DISCUSSION

Reactive oxygen species (ROS) are central signaling molecules in plant responses to abiotic stress, yet the contribution of distinct intracellular ROS sources and their coordination across organelles remain incompletely understood. Glycolate oxidase (GOX) is the first enzyme of the photorespiratory pathway allocated in peroxisomes. The production of hydrogen peroxide (H_2_O_2_) during the oxidation of glycolate to glyoxylate by GOX is a key feature of photorespiration (Foyer et al., 2009). Five genes annotated as glycolate oxidases are contained within the genome of the Arabidopsis plant playing GOX1 and GOX2 an essential role in basic metabolism as key enzyme within the photorespiration pathway (Foyer et al., 2009). In this study, we identify GOX1 and GOX2 as key modulators of redox homeostasis and mitochondrial function under control and cadmium (Cd) stress, revealing a previously underappreciated role for the photorespiratory pathway in interorganelle redox signaling.

Our data demonstrate that GOX1 and GOX2 fulfill non–redundant redox functions. GOX1 primarily sustains basal photorespiratory flux and regulates steady–state redox balance, whereas GOX2 is essential for stress–induced redox signaling and transcriptional reprogramming under Cd stress. This functional divergence aligns with previous findings showing that loss of GOX1, but not GOX2, suppresses the oxidative phenotype of catalase–deficient (*cat2*) plants and attenuated the SA responses in *cat2*, establishing GOX1 as the dominant metabolic source of photorespiratory H_2_O_2_ (Kerchev et al., 2016; Yang et al., 2025). Consistently, we observe that mitochondria in *gox1* plants exhibit a more oxidized redox state even under non–stress conditions, indicating that GOX1 contributes to basal redox tone across organelle boundaries.

In contrast, the most striking feature of our transcriptomic analysis is the strong dependence of Cd–responsive gene expression on GOX2. A large fraction of genes regulated by Cd in WT plants, including those associated with mitochondrial electron transport, antioxidant metabolism, and detoxification, fail to be regulated in *gox2* mutants. This places GOX2 upstream of transcriptional programs linked to redox acclimation and highlights its role as a signaling–competent ROS source, rather than a purely metabolic enzyme. Such a distinction between ROS production for metabolism versus signaling is increasingly recognized as a defining feature of compartment–specific redox control. Our results point to mitochondria as a central integration point of GOX–dependent redox regulation. Thus, beyond carbon flux, both organelles are major intracellular sources of reactive oxygen species and engage in bidirectional redox communication. Peroxisome-derived H_2_O_2_ acts as an early redox signal that influences alternative oxidase (AOX) activity, and respiratory flexibility, while mitochondrial redox status feeds back to regulate peroxisomal ROS metabolism and antioxidant capacity (Mhamdi et al., 2012; Noctor et al., 2002).

Cd exposure in WT plants induces respiratory activity alongside transcriptional up–regulation of electron transport chain components and the alternative oxidase AOX. The activation of AOX in response to Cd has been reported by Keunen et al. (2013) who suggested that it could limit Cd-dependent ROS production in these organelles. Finger-Teixeira et al. (2021) also have reported in soybean roots a Cd-dependent increased the respiration in the complex IV, while the heavy metal reduced the respiration in the complex III, and abolished the respiratory control of isolated mitochondria, especially when L-malate was used as substrate. In our study, Cd-dependent changes were attenuated or mis–regulated in both *gox1* and *gox2* mutants, indicating that photorespiratory redox signals are required to reprogram mitochondrial respiration under metal stress. Notably, despite minimal changes in mitochondrial redox status in Cd–treated WT plants, both GOX mutants display constitutively elevated mitochondrial oxidation, underscoring the role of GOX–derived signals in maintaining mitochondrial redox homeostasis rather than simply responding to stress–induced ROS accumulation. The absence of changes in the redox state of the mitochondria in WT plants due to Cd treatment could be due to the activation of detoxification mechanisms, mainly related with GST activities, and mitochondria-nucleus signalling mechanisms observed in the transcriptome. These mechanisms may already be activated in the mutants given that mitochondrial metabolism is altered in these lines. Thus, although energetically demanding, photorespiration is essential for a proper photosynthesis, carbon allocation, nitrogen and sulfur assimilation (Bauwe et al., 2012; Dellero et al., 2016; Florian et al., 2013; Timm et al., 2025). Beyond the metabolic role of photorespiration, GOX–derived ROS have been implicated in diverse physiological and defense processes, including innate immunity, hormone biosynthesis and senescence (Dellero et al., 2016; Rojas et al., 2012), as well as in response to salinity, regulating the uptake and translocation of Na^+^ and Cl^-^ (Benslima et al., 2026).

The coordination between peroxisomal H_2_O_2_ production and mitochondrial function supports the concept of a redox–signaling axis linking peroxisomes and mitochondria. Recent studies have shown that the glycolate oxidase–catalase (GC) module operates as a dynamic redox switch, modulating peroxisomal H_2_O_2_ levels in response to Ca²⁺–dependent systemic signals (Li et al., 2023). Our findings extend this framework by demonstrating that GOX–dependent redox signals propagate to mitochondria and shape respiratory and transcriptional responses under Cd stress (Fig. 7). These observations reinforce the view that photorespiratory H_2_O_2_ functions as a context–dependent signaling molecule, whose effects depend on enzymatic source, compartmentalization, and downstream redox relay mechanisms (Timm et al., 2025). In this regard, our data position GOX2 as a critical node coupling peroxisomal ROS production to nuclear gene expression and mitochondrial adaptation (Fig. 7). However, the precise mechanisms linking these signals to mitochondrial transcriptional regulation remain to be elucidated. Mitochondria and peroxisomes are highly dynamic organelles in plant cells, and their physical and functional interactions constitute an important layer of metabolic regulation. Rather than acting in isolation, these compartments communicate through membrane contact sites, metabolic exchange, and coordinated signaling pathways. In photosynthetic tissues, both organelles are frequently found in close proximity to each other and to chloroplasts, facilitating processes such as photorespiration.

**Figure 7:**
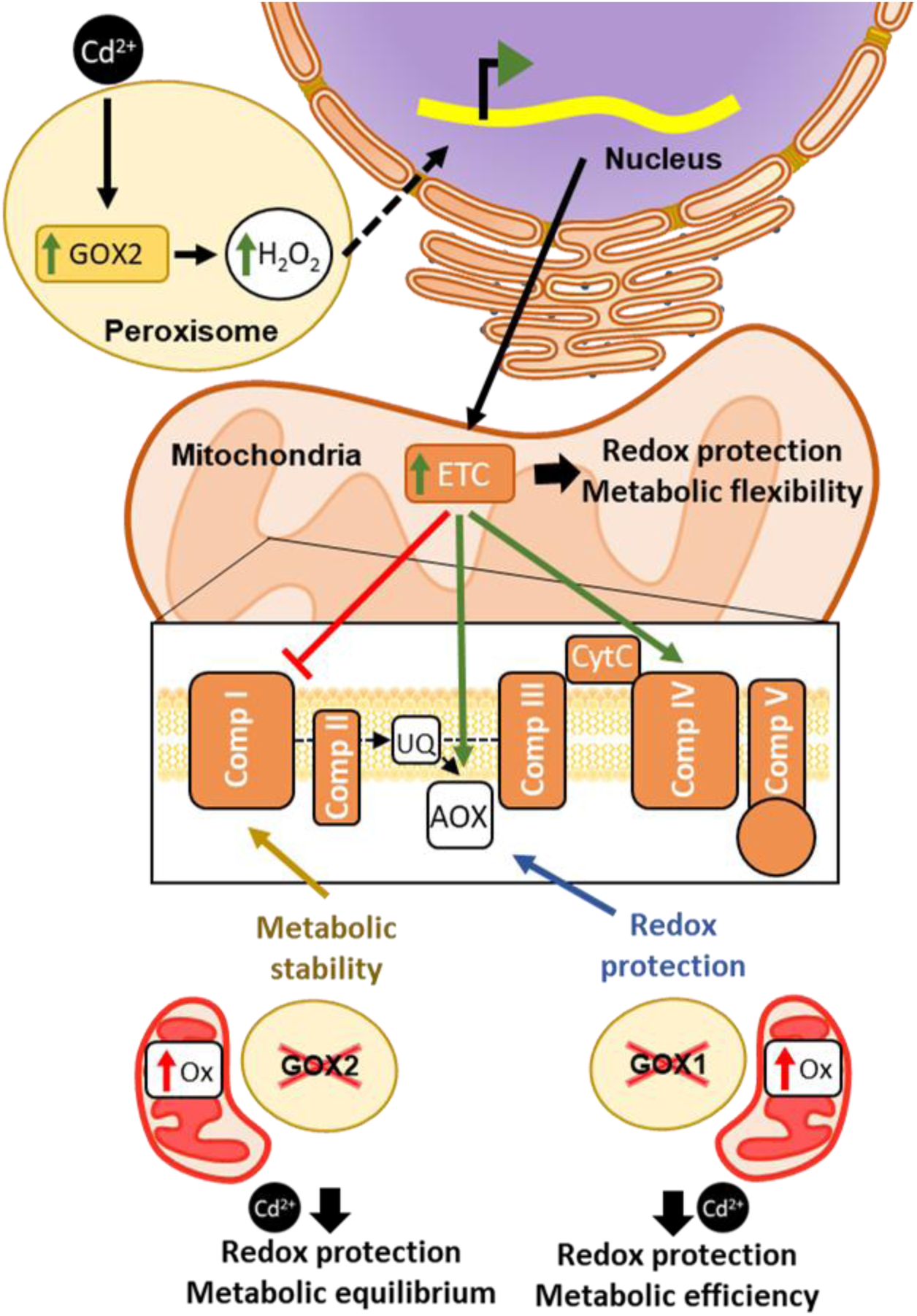
GOX1 and GOX2 modulate redox state of mitochondria and function under control and cadmium (Cd) stress. Treatment of Arabidopsis plants with cadmium (Cd) induces peroxisomal glycerol-3-phosphate oxidase (GOX) activity, which is associated with increased GOX2-dependent hydrogen peroxide (H_2_O_2_) production and transcriptional changes specific to this isoform. Some of these transcriptional changes reprogram the mitochondrial electron transport chain, inhibiting Complex I and inducing Complex IV and AOX, prioritizing mitochondrial redox protection and stress tolerance over maximal energy efficiency. Under control conditions however, the absence of the isoform GOX1 (*gox1*) produced a strong oxidation of mitochondria and an increase in mitochondrial AOX activity that promotes respiratory flexibility, limits excessive ROS formation, and enables redox homeostasis at the expense of ATP yield, thereby supporting stress acclimation rather than maximal energy efficiency. Under Cd stress this mutant maintain AOX activation and increase Complex I, improving metabolic efficiency. The absence of the isoform GOX2 (*gox2*) under control conditions also showed a higher oxidative state than in WT. These mutants showed an induced Complex I and, in less extent, AOX, supporting high respiratory capacity while reinforcing mitochondrial redox homeostasis and increasing metabolic stability. Under Cd stress, this mutant maintains AOX and reach a metabolic equilibrium between Complex I and Complex IV.

In conclusion, this work identifies photorespiratory glycolate oxidases as central components of plant redox signaling networks. We propose a model in which GOX1 maintains basal redox equilibrium, while GOX2 enables stress–induced redox signaling that coordinates mitochondrial function and transcriptional reprogramming during Cd exposure (Fig. 7). These findings redefine photorespiration as an active participant in redox communication rather than a collateral metabolic pathway and highlight the peroxisome–mitochondria interface as a key regulatory hub in cellular redox homeostasis.

## ACKNOWLEDGEMENTS

This study was co-funded by the Spanish Ministry of Science, Innovation and Universities, the “Agencia Estatal de Investigación” and the European Regional Development Fund (AEI/MICIN/ERDF; grants PID2021-122280NB-I00 and PID2024-155296NB-I00). P.R-R was supported by Grant A-EXP-161-UGR23 funded by Consejería de Universidad, Investigación e Innovación and by ERDF Andalusia Program 2021-2027, and by Grant RYC2023-043758-I funded by MICIU/AEI/10.13039/501100011033 and by ESF+. MAP-V was supported by MCIU Research Personnel Training (FPI) grant BES-2016-076518.

## CRediT authorship contribution statement

**AM Collado-Arenal**: Investigation, Formal analysis, Validation, Writing – review & editing. **M Rodríguez-Serrano**: Investigation, Formal analysis, Validation, Writing – review & editing. **MA Peláez-Vico**: Investigation, Formal analysis, Validation, Writing – review & editing. **LC Terrón-Camero**: Investigation, Software, Formal analysis, Writing – review & editing. **FL Pérez-Gordillo**: Formal analysis, Validation, Writing – review & editing. **P Ranea-Robles**: Supervision, Writing – review & editing. **LC López**: Writing – review & editing. **LM Sandalio**: Funding acquisition, Resources, Supervision, Contextualization, Writing – review & editing. **MC Romero-Puertas**: Writing – review & editing, Writing – original draft, Supervision, Project administration, Investigation, Funding acquisition, Conceptualization, Formal analysis, Validation.

## Data availability statement

The datasets presented in this study can be found in online repositories. The names of the repository/repositories and accession number(s) can be found at: https://www.ncbi.nlm.nih.gov/genbank/, GSE199325.

## REFERENCE LIST

1. Aebi H. Catalase in Vitro. Methods Enzymol 1984;105:121–6. 10.1016/S0076-6879(84)05016-3,.

2. Bauwe H, Hagemann M, Kern R, Timm S. Photorespiration has a dual origin and manifold links to central metabolism. Curr Opin Plant Biol 2012;15:269–75. 10.1016/j.pbi.2012.01.008.

3. Benslima W, Hafsi C, Espinosa J, Yun P, Romero-Puertas MC, Shabala S, et al. The role of glycolate oxidase in regulating Arabidopsis thaliana response to short-term salt stress. Plant Physiol Biochem PPB 2026;232:111159. 10.1016/j.plaphy.2026.111159.

4. Collado-Arenal AM, Exposito-Rodriguez M, Mullineaux PM, Olmedilla A, Romero-Puertas MC, Sandalio LM. Cadmium exposure induced light/dark- and time-dependent redox changes at subcellular level in Arabidopsis plants. J Hazard Mater 2024;477. 10.1016/J.JHAZMAT.2024.135164.

5. Cuypers A, Vanbuel I, Iven V, Kunnen K, Vandionant S, Huybrechts M, et al. Cadmium-induced oxidative stress responses and acclimation in plants require fine-tuning of redox biology at subcellular level. Free Radic Biol Med 2023;199:81–96. 10.1016/j.freeradbiomed.2023.02.010.

6. Dellero Y, Jossier M, Glab N, Oury C, Tcherkez G, Hodges M. Decreased glycolate oxidase activity leads to altered carbon allocation and leaf senescence after a transfer from high CO2 to ambient air in Arabidopsis thaliana. J Exp Bot 2016;67:3149–63. 10.1093/JXB/ERW054.

7. Finger-Teixeira A, Ishii-Iwamoto EL, Marchiosi R, Coelho ÉMP, Constantin RP, dos Santos WD, et al. Cadmium uncouples mitochondrial oxidative phosphorylation and induces oxidative cellular stress in soybean roots. Environ Sci Pollut Res 2021;28:67711–23. 10.1007/S11356-021-15368-2/FIGURES/7.

8. Florian A, Araújo WL, Fernie AR. New insights into photorespiration obtained from metabolomics. Plant Biol 2013;15:656–66. 10.1111/J.1438-8677.2012.00704.X;PAGEGROUP:STRING:PUBLICATION.

9. Food and Agriculture Organization of the United Nations. Saving our soils by all earthly ways possible 2022.

10. Foyer CH, Bloom AJ, Queval G, Noctor G. Photorespiratory metabolism: Genes, mutants, energetics, and redox signaling. Annu Rev Plant Biol 2009;60:455–84. 10.1146/ANNUREV.ARPLANT.043008.091948/1.

11. Foyer CH, Noctor G. Redox Homeostasis and Signaling in a Higher-CO2 World. Annu Rev Plant Biol 2020;71:157–82. 10.1146/ANNUREV-ARPLANT-050718-095955.

12. Fu Z, Xi S. The effects of heavy metals on human metabolism. Toxicol Mech Methods 2020;30:167–76. 10.1080/15376516.2019.1701594.

13. Garmier M, Carroll AJ, Delannoy E, Vallet C, Day DA, Small ID, et al. Complex I dysfunction redirects cellular and mitochondrial metabolism in Arabidopsis. Plant Physiol 2008;148:1324–41. 10.1104/PP.108.125880.

14. Garnier L, Simon-Plas F, Thuleau P, Agnel JP, Blein JP, Ranjeva R, et al. Cadmium affects tobacco cells by a series of three waves of reactive oxygen species that contribute to cytotoxicity. Plant, Cell Environ 2006;29:1956–69. 10.1111/j.1365-3040.2006.01571.x.

15. Gupta DK, Pena LB, Romero-Puertas MC, Hernández A, Inouhe M, Sandalio LM. NADPH oxidases differentially regulate ROS metabolism and nutrient uptake under cadmium toxicity. Plant Cell Environ 2017;40:509–26. 10.1111/pce.12711.

16. Hafsi C, Collado-Arenal AM, Wang H, Sanz-Fernández M, Sahrawy M, Shabala S, et al. The role of NADPH oxidases in regulating leaf gas exchange and ion homeostasis in Arabidopsis plants under cadmium stress. J Hazard Mater 2022;429:128217. 10.1016/J.JHAZMAT.2022.128217.

17. Hou D, Jia X, Wang L, McGrath SP, Zhu YG, Hu Q, et al. Global soil pollution by toxic metals threatens agriculture and human health. Science 2025;388:316–21. 10.1126/SCIENCE.ADR5214/SUPPL_FILE/SCIENCE.ADR5214_MDAR_REPRODUCIBILITY_CHECKLIST.PDF.

18. Hu Z, Vanderhaeghen R, Cools T, Wang Y, De Clercq I, Leroux O, et al. Mitochondrial Defects Confer Tolerance against Cellulose Deficiency. Plant Cell 2016;28:2276. 10.1105/TPC.16.00540.

19. Kerchev P, Waszczak C, Lewandowska A, Willems P, Shapiguzov A, Li Z, et al. Lack of GLYCOLATE OXIDASE1, but not GLYCOLATE OXIDASE2, attenuates the photorespiratory phenotype of CATALASE2-deficient Arabidopsis. Plant Physiol 2016;171:1704–19. 10.1104/pp.16.00359.

20. Kerr MW, Groves D. Purification and properties of glycollate oxidase from Pisum sativum leaves. Phytochemistry 1975;14:359–62. 10.1016/0031-9422(75)85090-4.

21. Keunen E, Jozefczak M, Remans T, Vangronsveld J, Cuypers A. Alternative respiration as a primary defence during cadmium-induced mitochondrial oxidative challenge in Arabidopsis thaliana. Environ Exp Bot 2013;91:63–73. 10.1016/j.envexpbot.2013.02.008.

22. Li X, Chen L, Zeng Xiaoyue, Wu K, Huang J, Liao M, et al. Wounding induces a peroxisomal H2 O2 decrease via glycolate oxidase-catalase switch dependent on glutamate receptor-like channel-supported Ca2+ signaling in plants. Plant J 2023;116:1325–41. 10.1111/TPJ.16427.

23. Maliandi M V., Rius SP, Busi M V., Gomez-Casati DF. A simple method for the addition of rotenone in Arabidopsis thaliana leaves. Plant Signal Behav 2015;10:e1073871. 10.1080/15592324.2015.1073871.

24. Mhamdi A, Noctor G, Baker A. Plant catalases: Peroxisomal redox guardians. Arch Biochem Biophys 2012;525:181–94. 10.1016/J.ABB.2012.04.015.

25. Molina-Moya E, Rodríguez-González A, Peláez-Vico MA, Sandalio LM, Romero-Puertas MC. Peroxisomal dependent signalling and dynamics modulate plant stress responses: reactive oxygen and nitrogen species as key molecules. J Exp Bot 2025. 10.1093/JXB/ERAF072.

26. Murashige T, Skoog F. A revised medium for rapid growth and bio assays with tobacco tissue cultures. Physiol Plant 1962;15:473–97. 10.1111/J.1399-3054.1962.TB08052.X.

27. Noctor G, Veljovic-Jovanovic S, Driscoll S, Novitskaya L, Foyer CH. Drought and oxidative load in the leaves of C3 plants: a predominant role for photorespiration? Ann Bot 2002;89 Spec No:841–50. 10.1093/AOB/MCF096.

28. Okereafor U, Makhatha M, Mekuto L, Uche-Okereafor N, Sebola T, Mavumengwana V. Toxic Metal Implications on Agricultural Soils, Plants, Animals, Aquatic life and Human Health. Int J Environ Res Public Health 2020;17. 10.3390/IJERPH17072204.

29. Peco JD, Campos JA, Romero-Puertas MC, Olmedilla A, Higueras P, Sandalio LM. Characterization of mechanisms involved in tolerance and accumulation of Cd in Biscutella auriculata L. Ecotoxicol Environ Saf 2020;201:110784. 10.1016/j.ecoenv.2020.110784.

30. Pérez-Chaca MV, Rodríguez-Serrano M, Molina AS, Pedranzani HE, Zirulnik F, Sandalio LM, et al. Cadmium induces two waves of reactive oxygen species in Glycine max (L.) roots. Plant, Cell Environ 2014;37:1672–87. 10.1111/pce.12280.

31. Remans T, Keunen E, Bex GJ, Smeets K, Vangronsveld J, Cuypers A. Reliable gene expression analysis by reverse transcription-quantitative PCR: Reporting and minimizing the uncertainty in data accuracy. Plant Cell 2014;26:3829–37. 10.1105/TPC.114.130641.

32. Reumann S, Babujee L, Changle M, Wienkoop S, Siemsen T, Antonicelli GE, et al. Proteome analysis of Arabidopsis leaf peroxisomes reveals novel targeting peptides, metabolic pathways, and defense mechanisms. Plant Cell 2007;19:3170–93. 10.1105/TPC.107.050989.

33. Rodríguez-Serrano M, Romero-Puertas MC, Sanz-Fernández M, Hu J, Sandalio LM. Peroxisomes extend peroxules in a fast response to stress via a reactive oxygen species-mediated induction of the peroxin PEX11a. Plant Physiol 2016;171:1665–74. 10.1104/pp.16.00648.

34. Rojas CM, Senthil-Kumar M, Wang K, Ryu CM, Kaundal A, Mysore KS. Glycolate Oxidase Modulates Reactive Oxygen Species–Mediated Signal Transduction during Nonhost Resistance in Nicotiana benthamiana and Arabidopsis. Plant Cell 2012;24:336–52. 10.1105/TPC.111.093245.

35. Romero-Puertas MC, McCarthy I, Sandalio LM, Palma JM, Corpas FJ, Gómez M, et al. Cadmium toxicity and oxidative metabolism of pea leaf peroxisomes. Free Radic Res 1999;31 Suppl. 10.1080/10715769900301281.

36. Romero-Puertas MC, Rodríguez-Serrano M, Corpas FJ, Gómez M, Del Río LA, Sandalio LM. Cadmium-induced subcellular accumulation of O2·− and H2O2 in pea leaves. Plant Cell Environ 2004;27:1122–34. 10.1111/J.1365-3040.2004.01217.X.

37. Rosenwasser S, Fluhr R, Joshi JR, Leviatan N, Sela N, Hetzroni A, et al. ROSMETER: A Bioinformatic Tool for the Identification of Transcriptomic Imprints Related to Reactive Oxygen Species Type and Origin Provides New Insights into Stress Responses. Plant Physiol 2013;163:1071–83. 10.1104/PP.113.218206.

38. Sandalio LM, Collado-Arenal AM, Romero-Puertas MC. Deciphering peroxisomal reactive species interactome and redox signalling networks. Free Radic Biol Med 2023;197:58–70. 10.1016/j.freeradbiomed.2023.01.014.

39. Sew YS, Ströher E, Holzmann C, Huang S, Taylor NL, Jordana X, et al. Multiplex micro-respiratory measurements of Arabidopsis tissues. New Phytol 2013;200:922–32. 10.1111/NPH.12394.

40. Terrón-Camero LC, Peláez-Vico MÁ, Rodríguez-González A, del Val C, Sandalio LM, Romero-Puertas MC. Gene network downstream plant stress response modulated by peroxisomal H2O2. Front Plant Sci 2022;13. 10.3389/FPLS.2022.930721.

41. Terrón-Camero LC, Rodríguez-Serrano M, Sandalio LM, Romero-Puertas MC. Nitric oxide is essential for cadmium-induced peroxule formation and peroxisome proliferation. Plant Cell Environ 2020;43:2492–507. 10.1111/PCE.13855.

42. Timm S, Sun H, Hagemann M, Huang W, Fernie AR. An old dog with new tricks-the value of photorespiration as a central metabolic hub with implications for environmental acclimation. Plant Physiol 2025;198. 10.1093/PLPHYS/KIAF258.

43. Wrzaczek M, Brosché M, Salojärvi J, Kangasjärvi S, Idänheimo N, Mersmann S, et al. Transcriptional regulation of the CRK/DUF26 group of receptor-like protein kinases by ozone and plant hormones in Arabidopsis. BMC Plant Biol 2010;10. 10.1186/1471-2229-10-95.

44. Yan H, Peng Z, Zhang H, Wang B, Xu W, He Z. Cadmium Minimization in Crops: A Trade-Off With Mineral Nutrients in Safe Breeding. Plant Cell Environ 2025;48:838–51. 10.1111/PCE.15182.

45. Yang T, Mu X, Yu M, Ergashev U, Zhu Y, Shi N, et al. Consecutive oxidative stress in CATALASE2-deficient Arabidopsis negatively regulates Glycolate Oxidase1 activity through S-nitrosylation. Physiol Plant 2025;177. 10.1111/PPL.70040.

