## Supplemental Figures S1-S3 for "The lack of peroxisomal Glycolate Oxidases 1 and 2 influences mitochondrial electron transport chain and its redox state under control and cadmium stress"

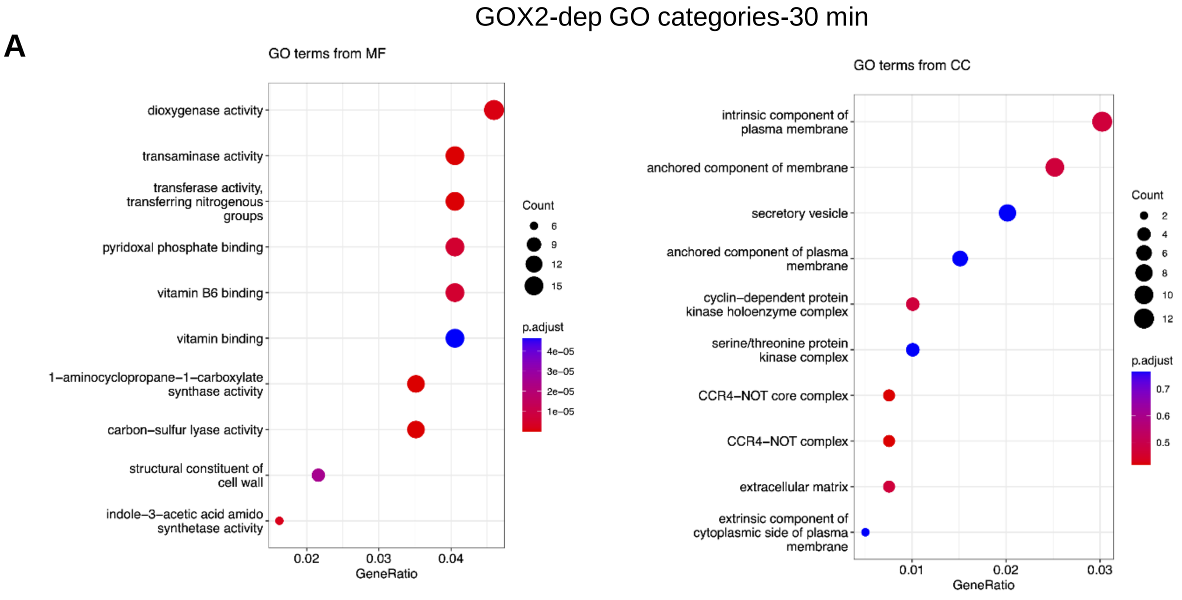


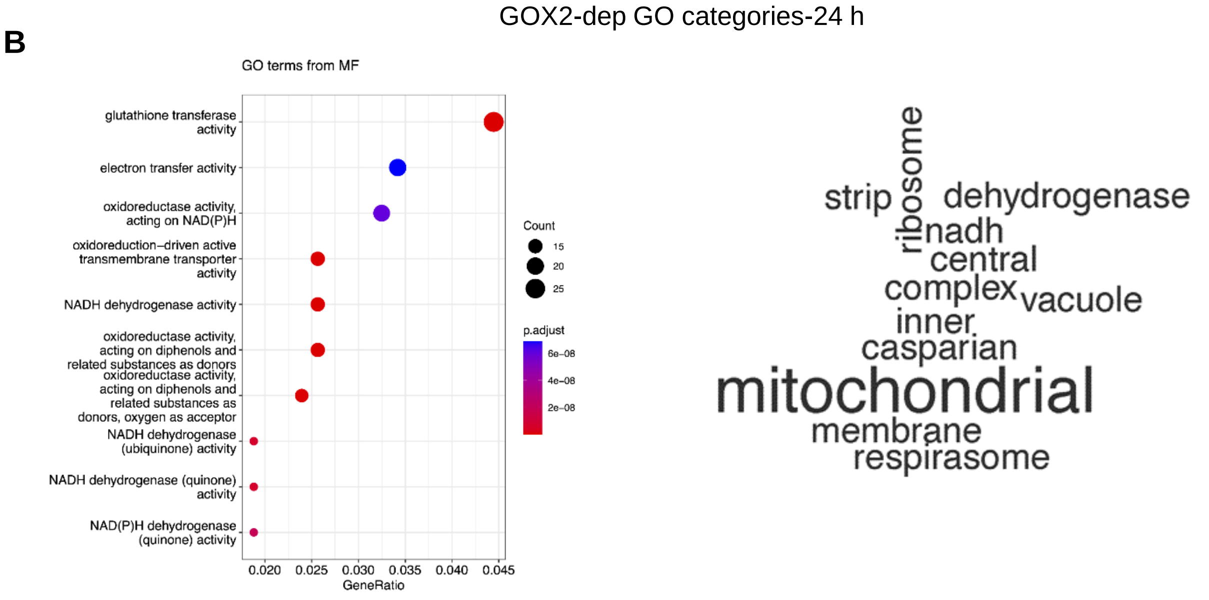


**Suppl. Fig. S1. Main Molecular Function and Cellular Components in GOX2-dependent genes in response to Cd treatment.** Main Molecular Function and Cellular Components categories after gene ontology (GO) enrichment of GOX2-dependent genes at 30min (A) and 24 h (B) after Cd treatment (Figure 3; Suppl. Table S3 and S4). Normed to frequency of class over all ID numbers on *x* axes.

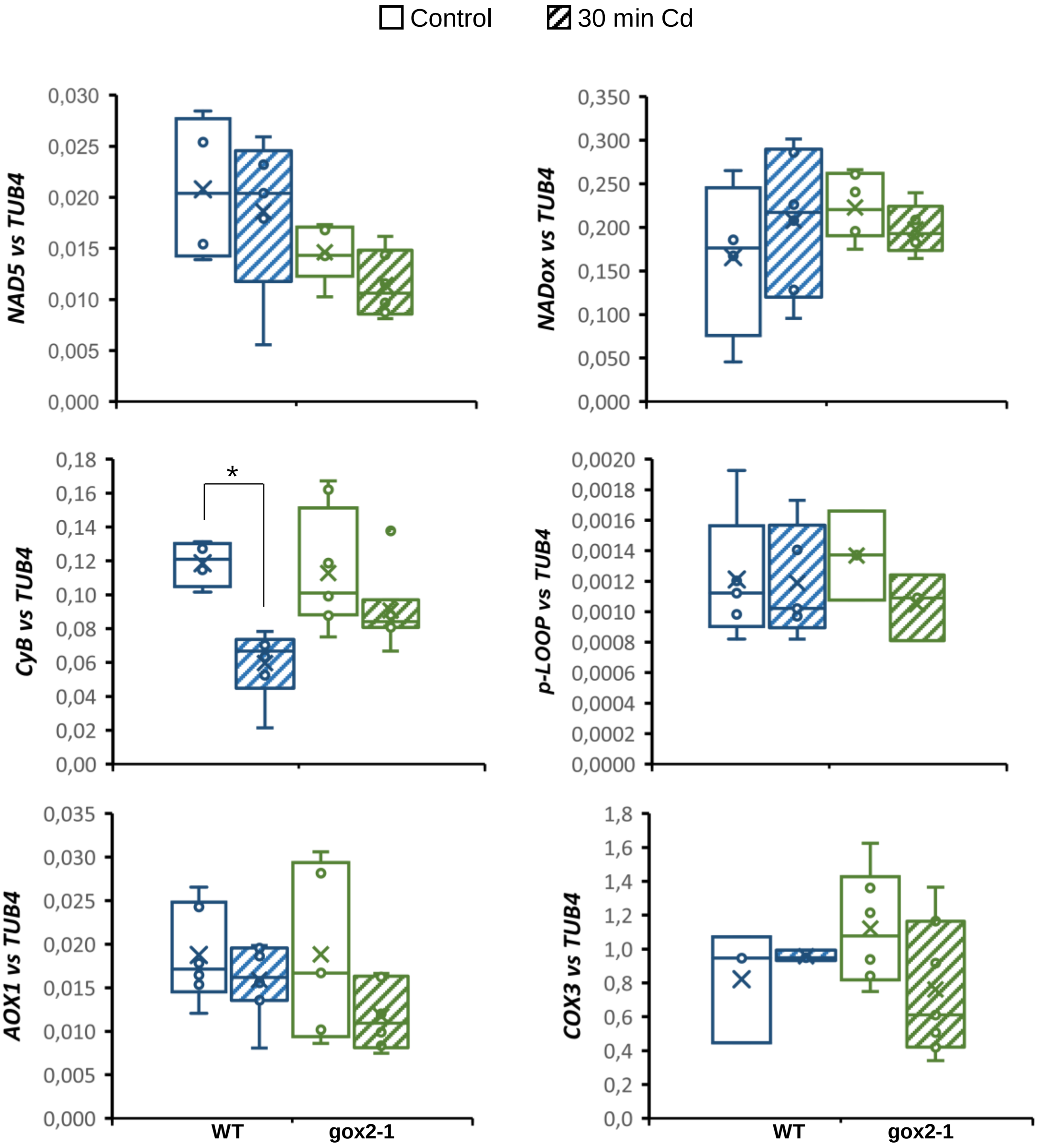


**Suppl. Fig. S2: Early transcriptomic changes induced by Cd in mitochondrial ETC genes in WT and *gox2* mutant.** Quantitative real-time PCR analyses of genes related to ETC (Table 1) in WT and *gox2* seedlings treated with Cd for 30 min. Each gene was normalized against *TUB4* expression. Primers used are described in Suppl. Table S1. Asterisks denote significant differences between data according to the Student's t-test ($p$-value $< 0.05$). The absence of an asterisk denotes no significant differences.

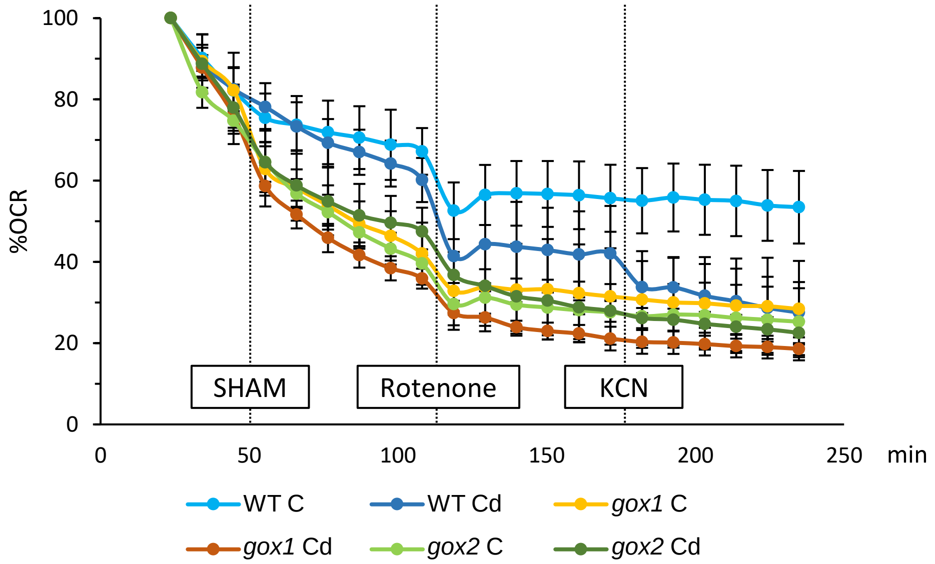


**Supplemental Figure S3: Cadmium effect on mitochondrial %OCR in WT and *gox* mutants.** Sequential profile of mitochondrial respiration in leave discs from WT and *gox1* and *gox2* mutants treated (Cd) or not (C) with Cd for 24 h. The OCR was measured using an Agilent Seahorse XFe24 and inhibitors effect: SHAM (alternative oxidase inhibitor); rotenone (Complex I inhibitor); and KCN (Complex IV inhibitor).
